# A 5’ UTR Language Model for Decoding Untranslated Regions of mRNA and Function Predictions

**DOI:** 10.1101/2023.10.11.561938

**Authors:** Yanyi Chu, Dan Yu, Yupeng Li, Kaixuan Huang, Yue Shen, Le Cong, Jason Zhang, Mengdi Wang

**Affiliations:** Department of Pathology, Stanford University School of Medicine, Stanford, CA 94305, USA; RVAC Medicines, Waltham, MA 02451, USA; Department of Electrical and Computer Engineering, Princeton University, Princeton, NJ 08544, USA; Center for Statistics and Machine Learning, Princeton University, Princeton, NJ 08544, USA

**Keywords:** 5’ untranslated region, mRNA, language model, semi-supervised learning, ribosome loading, translation efficiency, gene expression, internal ribosome entry site

## Abstract

The 5’ UTR, a regulatory region at the beginning of an mRNA molecule, plays a crucial role in regulating the translation process and impacts the protein expression level. Language models have showcased their effectiveness in decoding the functions of protein and genome sequences. Here, we introduced a language model for 5’ UTR, which we refer to as the UTR-LM. The UTR-LM is pre-trained on endogenous 5’ UTRs from multiple species and is further augmented with supervised information including secondary structure and minimum free energy. We fine-tuned the UTR-LM in a variety of downstream tasks. The model outperformed the best-known benchmark by up to 42% for predicting the Mean Ribosome Loading, and by up to 60% for predicting the Translation Efficiency and the mRNA Expression Level. The model also applies to identifying unannotated Internal Ribosome Entry Sites within the untranslated region and improves the AUPR from 0.37 to 0.52 compared to the best baseline. Further, we designed a library of 211 novel 5’ UTRs with high predicted values of translation efficiency and evaluated them via a wet-lab assay. Experiment results confirmed that our top designs achieved a 32.5% increase in protein production level relative to well-established 5’ UTR optimized for therapeutics.

## 1. Introduction

The 5’ untranslated region (5’ UTR) is a region at the beginning of an mRNA that precedes the coding sequence of the protein. It plays a critical role in regulating the translation from mRNA to proteins, as it can influence the stability, localization, and translation of the mRNA molecule^1^. There has been a significant amount of research^2–8^ exploring the role of the 5’ UTR in various biological processes, including its secondary structure^2^, RNA-binding proteins that may interact with it^3^, and the effect of mutations within the 5’ UTR on the gene expression^4^. The complex functions of mRNA and their potential implications for human health and diseases underscore the necessity for more universally applicable computational approaches.

Investigation into the role of 5’ UTRs encompasses various aspects of translational control. With the growing interest in studying and designing 5’ UTRs, various computational tools^5–9, 7, 8^ have been developed to study its functions. For example, the ribosome load measures the number of ribosomes engaged in translating a given mRNA at a given time. Supervised machine learning models were shown to predict the mean ribosome loading (MRL)^6, 7, 9^ based on the UTR sequence or its biological features. Additionally, RNABERT^10^ and RNA-FM^9^ are language models specific to RNA sequences and were shown useful for predicting the MRL. Further, 5’ UTR is also predictive of the mRNA translation efficiency (TE)^5, 8^ that quantifies the rate of translation into proteins and the mRNA expression level (EL)^8^ that reflects the relative abundance of the mRNA transcript in the cell. While there exist specialized machine learning models for individual tasks, there lacks a methodology to decode these functions of 5’ UTRs in a unified way.

In this study, we adopted the principled approach of a language model in order to extract meaningful semantic representations from untranslated regions of mRNA and further map them to predict functions of interest. Specifically, we developed a semi-supervised language model, which we refer to as the UTR-LM, trained using sequences of 5’ UTR from multiple data sources (Fig. 1). The transformer-based model is pre-trained to extract feature representations from the raw sequences via nucleotides masking and reconstruction. It also incorporated supervised information such as the secondary structure (SS) and the minimum free energy (MFE). We applied the UTR-LM and fine-tuned it for a variety of downstream tasks, such as predicting the mean ribosome loading, the mRNA translation efficiency, and the mRNA expression level. Experiment results showed that the UTR-LM accurately predicts these regulatory functions. When compared with existing baselines in each downstream task^6–11^, the UTR-LM demonstrated state-of-the-art performances across modalities and test sets. In particular, the UTR-LM outperforms the RNAFM^9^ by 42% for MRL prediction, and outperforms the FramePool^6^ by up to 60% for TE and EL predictions, in terms of the Spearman R score. The model generalized well to unseen data, especially human 5’ UTRs with varying lengths. Additionally, we adapted the UTR-LM to identify unannotated internal ribosome entry sites (IRESs)^11–14^, which are specific sequences within some mRNAs that enable ribosomes to initiate translation internally, bypassing the traditional cap-dependent mechanism. The UTR-LM outperforms IRESpy^11^ by 0.15 in terms of the AUPR score.

**Fig. 1.**
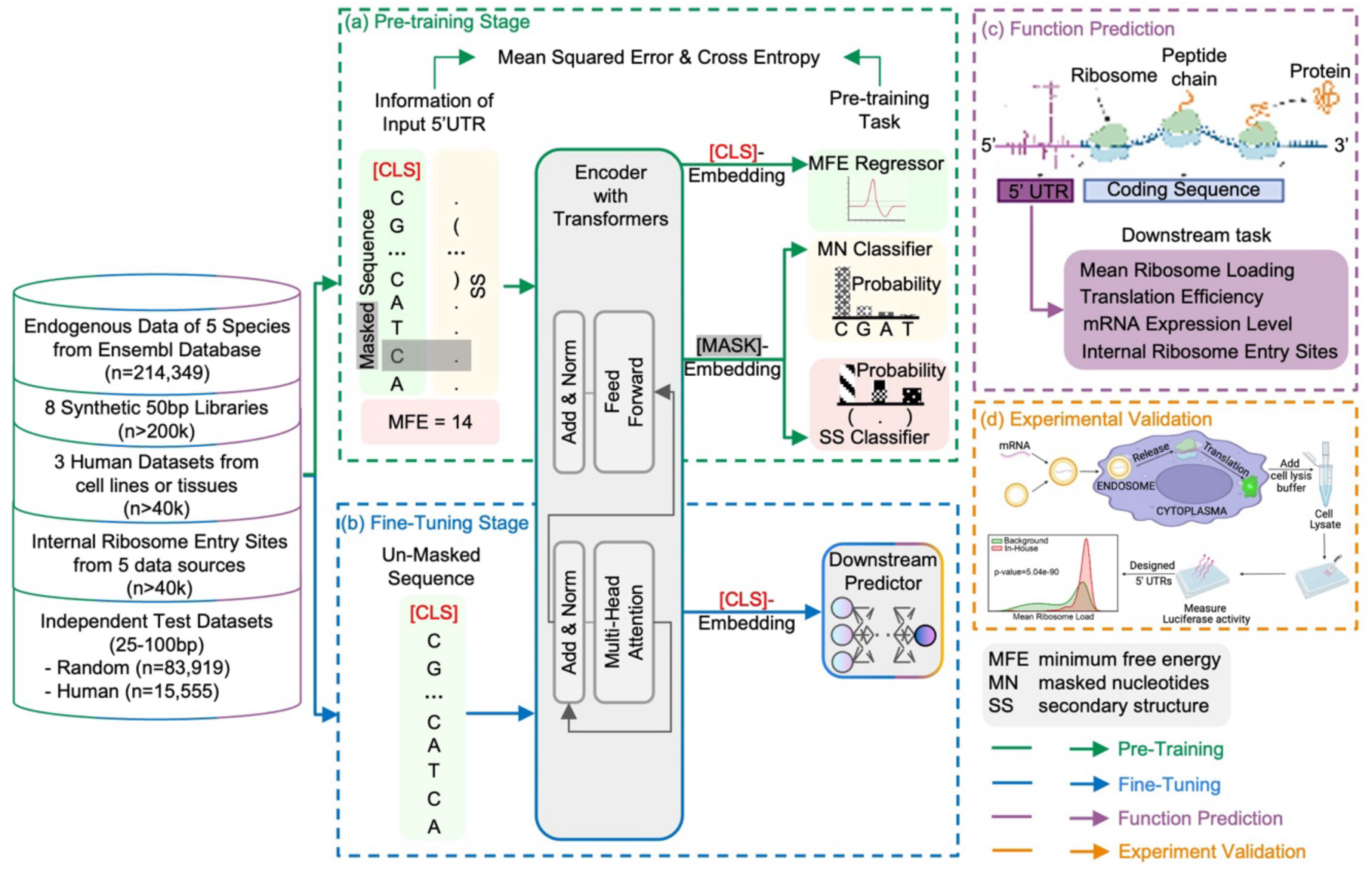
Overview of the UTR-LM model for 5’ UTR function prediction and design. (a) The input of the proposed pre-trained model is the 5’ UTR sequence, which is fed into the transformer layer through a randomly generated 128-dimensional embedding for each nucleotide and a special [CLS] token. The pre-training phase employs a combination of masked nucleotide (MN) prediction, 5’ UTR secondary structure (SS) prediction, and 5’ UTR minimum free energy (MFE) prediction. (b) Following pre-training, the [CLS] token is used for downstream task-specific training. (c) The UTR-LM is fine-tuned for downstream tasks such as predicting mean ribosome loading, translation efficiency, mRNA expression level, and internal ribosome entry site. (d) Designing an in-house library of 5’ UTRs with highly predicted translation efficiency and the wet-lab experimental validation employing mRNA transfection and luciferase assays. Drawings were created using BioRender.com.

Given the vital role of untranslated regions of mRNA in the translation process, artificially designing the 5’ UTR holds the potential to improve translation efficiency and optimize protein production^7^. Leveraging this biological principle, we designed an in-house library of 211 novel 5’ UTRs with high predicted values of translation efficiency. We conducted experiments of mRNA transfection and luciferase assay to evaluate our designs. Wet-lab experiments revealed that the top 5’ UTRs in our design library achieved up to 32.5% increase in protein production level, when compared to the benchmark NCA-7d-5’UTR^15^, which was optimized for encoding SARS-CoV-2 antigens and elicited strong immunity when delivered using liquid nanoparticle (LNP) in an in vivo vaccination experiment. The observed increase in protein production level suggests that the newly designed 5’ UTRs could have practical applications in various biotechnological processes. Further, we used the in-house data as an independent set to test UTR-LM for zero-shot fitness prediction, and showed that the model significantly outperformed other methods by up to 51% in terms of the Spearman R score.

Additionally, we analyzed the pre-trained embedding and the attention score of the UTR-LM. The embedding of UTR-LM is able to differentiate between species and capture features like the MFE. We also sought to detect motif patterns based on attention scores. In particular, the presence of the Kozak consensus sequence with higher GC content was found to be significant, a finding that aligns with previous research^16, 17^.

In summary, our study presents the UTR-LM, a novel semi-supervised language model for studying untranslated regions of mRNA and decoding its functions. This research holds promising implications for advancing our understanding of gene regulation and innovating therapeutic interventions.

## 2. Results

### 2.1. UTR-LM predicts the Mean Ribosome Loading accurately and generalizes to human 5’ UTRs

Ribosome loading refers to the number of ribosomes that are actively translating a specific mRNA molecule at any given time. It is a measure of how efficiently a particular mRNA is being translated into protein, and can be measured by various experimental techniques such as Ribo-seq or polysome profiling^18^. The ribosome loading can influence the rate of protein production and is influenced by factors such as the 5’ untranslated region (5’ UTR) sequence, secondary structures within the mRNA, and the availability of ribosomes. Designing the 5’ UTR can impact ribosome loading, which in turn can be utilized to optimize protein expression levels for various applications, including biotechnology and therapeutic protein production. Scientists had attempted to predict the effect of 5’ UTR sequence on the mean ribosome loading (MRL). Several machine learning models were developed for this specific task, including Optimus^7^ and FramePool^6^. RNA language models namely RNABERT^10^ and RNA-FM^9^ were also tested using this task.

In our study, we used the pre-trained foundation model for 5’ UTR and further fine-tuned it for the task of MRL prediction via semi-supervised learning. The baseline model is pre-trained on unlabeled 5’ UTR sequences from five species within the Ensembl database, employing the masked nucleotides (MN) task, and subsequently fine-tuned for the downstream MRL task prediction. As illustrated in Fig. 2a, we tested variants of the model with additional training, including raw sequences from the downstream library, supervised information such as secondary structure (SS), minimum free energy (MFE) and other biology features. Fig. 2b reveals that the baseline model alone attains satisfactory performance, and minor differences are observed among UTR-LM variants. For the final model, we chose the version that incorporates the downstream library, SS and MFE, and we will refer to it as UTR-LM MRL in the rest of the subsection.

**Fig. 2.**
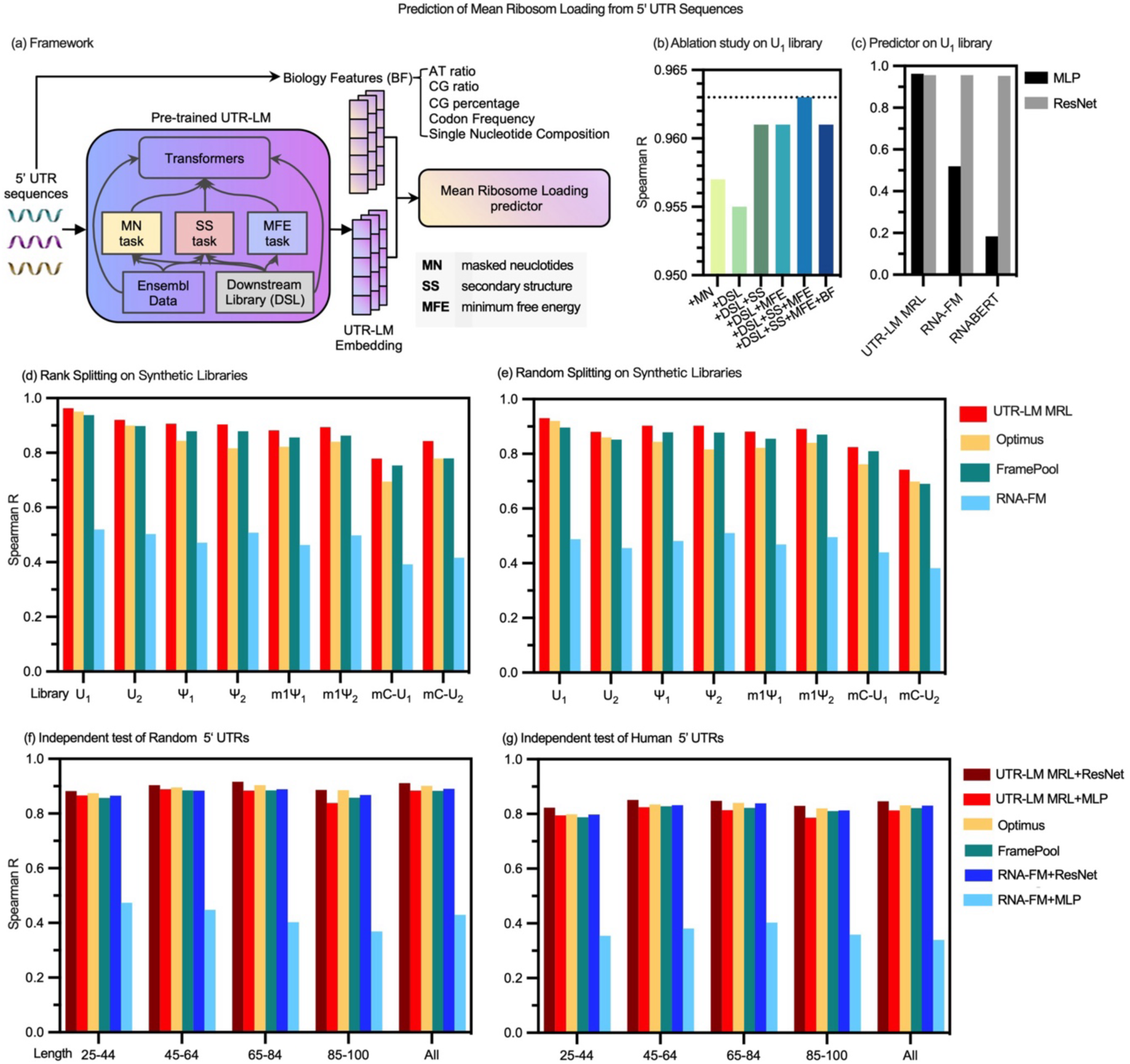
Prediction of mean ribosome loading based on 5’ UTR sequences. (a) Illustration of the UTR-LM framework, which includes variants integrating downstream library (DSL), secondary structure (SS), and minimum free energy (MFE) tasks during pre-training, along with biological features (BF) during downstream prediction. (b) Ablation study of UTR-LM hyperparameters under the U_1_ library (a sub-library of synthetic 50-nucleotide 5’ UTRs, see the Methods Section for details) with Rank Splitting. For subsequent experiments, we used the baseline UTR-LM enhanced by DSL, SS, and MFE, termed UTR-LM MRL. (c) Comparison of various pre-trained methods followed by either a simple multilayer perceptron (MLP) with 1 hidden layer or a complex 32-layer residual network (ResNet) under U_1_ library with Rank Splitting. (d-e) Evaluation of various methods across eight libraries with random 50-nucleotide 5’ UTRs with Rank Splitting and Random Splitting, respectively. (f-g) Evaluation of various methods using independent tests. In independent tests, we fine-tuned UTR-LM MRL and retrained baselines on 76,319 random 5’ UTRs (25-100 bp), and tested models on 7,600 random and 7,600 human 5’ UTRs.

We tested the UTR-LM MRL model with two downstream predictors: a simple multilayer perceptron (MLP) with 1 hidden layer and a much more complex 32-layer residual network (ResNet). In Fig. 2c, UTR-LM MRL followed by both predictors achieve similar performances, outperforming two other RNA foundation language models, i.e., RNA-FM and RNABERT, on this same task. Moreover, RNA-FM and RNABERT worked well with a deep ResNet predictor, but performed poorly with the simpler MLP predictor. This observation hints that RNA-FM and RNABERT rely on deep neural networks to learn the sequence-to-MRL relation. In contrast, our pre-trained UTR-LM MRL model does not rely on such complex predictors and achieves robustly high performances, indicating that it has extracted better semantic embeddings from 5’ UTR sequences.

We compared the UTR-LM MRL model with benchmark methods from the MRL prediction literature^6, 7, 9^ across eight synthetic libraries. This analysis focused on the 50-bp segment within the 5’ UTRs of genes closest to the start codon. Variants within this region are believed to be subject to stronger negative selection, likely because they can have more immediate effects on the gene’s ability to produce proteins^19^. We tested two different splitting strategies (Supplementary Materials B.3): Rank Splitting^7^ selects the 5’ UTRs with the highest read counts as the test data and uses the rest for training; Random Splitting splits data randomly into training and test sets. As illustrated in Fig. 2d,e, UTR-LM MRL consistently shows higher performance compared to other methods across all these tests. In particular, the UTR-LM MRL outperforms Optimus by up to 9%, outperforms FramePool by up to 6%, and outperforms RNAFM by up to 42%, in terms of the Spearman R score.

We aimed to assess whether our model, specifically trained on 50-bp synthetic sequences, could predict the regulatory functions of human 5’ UTR sequences with varying lengths. While human 5’ UTR sequences can span from tens to thousands of nucleotides, a mere 13% of them are less than 50-bp long. We used two datasets that were originally proposed by Sample et al.^7^ and later as independent tests in several studies^6, 7, 9^. They allow evaluating the model’s adaptability from the training data (including only synthetic 5’ UTRs of 50-bp length) to human 5’ UTRs of varying lengths. These datasets encompass both synthetic 5’ UTRs and human 5’ UTRs, with their lengths ranging from 25 to 100 bp, and they do not overlap with our training data. By following the length-based held-out testing approach suggested by Optimus^7^, we fine-tuned the UTR-LM MRL model and retrained available baselines^6, 7, 9^ on the 76,319 random 5’ UTRs with 25 to 100 bp. Then, we tested and compared UTR-LM MRL and baselines on 7,600 random and 7,600 human 5’ UTRs. Further details can be found in the Methods Section and Supplementary Materials B.4. Our results are illustrated in Fig. 2f,g. They show that the UTR-LM MRL, when paired with a ResNet downstream predictor, outperforms all other methods in both tests, while UTR-LM MRL with a simpler MLP exhibits similarly competitive performance. In addition, we analyzed the prediction results across various sequence lengths. The results demonstrate that UTR-LM MRL can be effectively extended to both longer and shorter 5’ UTRs. Notably, our model shows the state-of-the-art performance on human 5’ UTRs (Fig. 2g). In particular, the UTR-LM MRL outperforms Optimus, FramePool, and RNAFM by about 3%, in terms of Spearman R score. It highlights the generalizability of UTR-LM MRL for decoding functions of endogenous 5’ UTRs.

### 2.2. UTR-LM predicts the Translation Efficiency and mRNA Expression Level accurately

Protein production involves two primary processes: transcription and translation. The level of protein expression is highly dependent on the mRNA expression level (EL) and also the translation efficiency (TE) of the transcripts^20, 21^. EL is measured based on the relative abundance of the mRNA transcript in the cell and is quantified using RNA-seq RPKM^8^, where RPKM denotes Reads Per Kilobase of transcript per Million mapped reads. On the other hand, the TE of a gene, reflecting the rate of mRNA translation into protein, is calculated by dividing the Ribo-seq RPKM (indicative of ribosomal footprints on the mRNA) by the RNA-seq RPKM^8^.

In this section, we applied the pre-trained UTR-LM and fine-tuned it for TE and EL prediction. We used three endogenous datasets^8^ gathered from human muscle tissue (Muscle), human prostate cancer cell line PC3 (PC3), and human embryonic kidney 293T cell line (HEK) for training and testing. For benchmarking, we compared our method with a random forest model based on 3000+ handcrafted biological features by Cao et al.^8^ (Cao-RF), and several other sequence-based models including Optimus^7^, FramePool^6^, RNABERT^10^, and RNA-FM^9^. Here note that the Optimus, FramePool, RNABERT, and RNA-FM were not developed for the mRNA TE and EL tasks, thus for comparison we retrained these models using TE and EL datasets.

For the prediction of TE, we conducted ablation studies using various UTR-LM variants. Unlike the MRL task (as shown in Fig. 2b), the TE task (illustrated in Fig. 3a,b,c) reveals notable performance discrepancy among different UTR-LM variants. When augmented with downstream library and MFE loss, the fine-tuned model demonstrates substantially higher performance across various datasets. We observed that the use of the downstream library alone or SS could lead to performance degradation, possibly due to the fact that we had to truncate long sequences in the model and it might have introduced noticeable bias. We also observed that the additional use of biological features only affects the performance very minorly, so we chose to not include it in the final model. In the rest of the section, we chose the UTR-LM enhanced by the downstream library and MFE for the tasks TE and EL prediction. We will refer to this variant as UTR-LM TE for brevity. We also adapted the UTR-LM TE architecture to predict EL, which we refer to as UTR-LM EL.

**Fig. 3.**
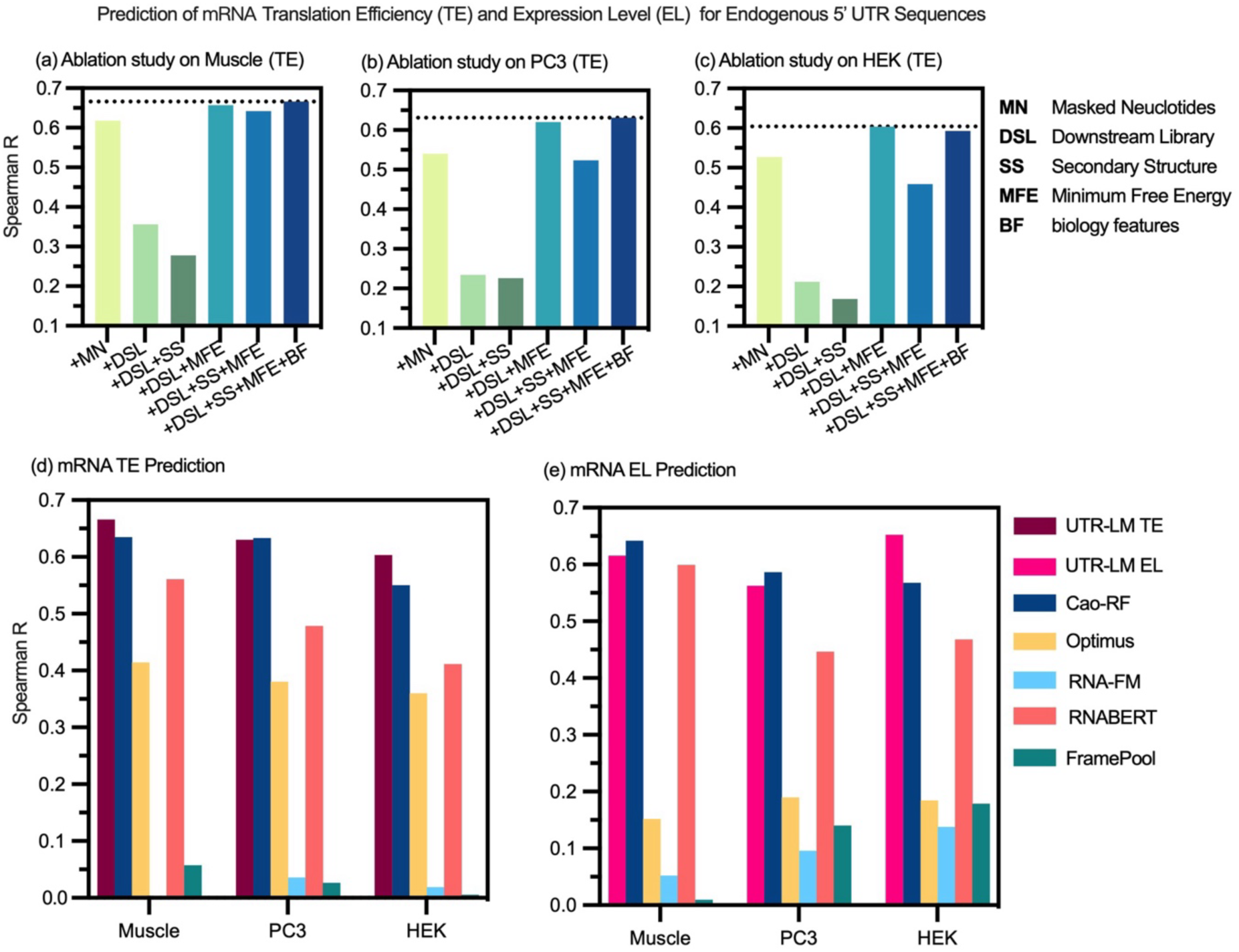
Prediction of mRNA translation efficiency (TE) and expression level (EL) for endogenous datasets. Sources for these datasets include human muscle tissue (Muscle), human prostate cancer cells (PC3), and human embryonic kidney 293T cells (HEK). (a-c) Ablation study of UTR-LM hyperparameters on TE tasks, including the downstream library’s 5’ UTRs (DSL), a downstream task-independent secondary structure (SS), and a downstream task-related minimum free energy (MFE) during the pre-training phase, and the inclusion of biological features (BF) during the downstream prediction phase. For the subsequent experiments, we used the baseline UTR-LM enhanced by DSL and MFE as the final model. (d) For the TE prediction, the UTR-LM model outperforms Cao-RF by up to 5% and outperforms Optimus by up to 25% in terms of Spearman R. (e) For the TE prediction, the UTR-LM model outperforms Cao-RF by up to 8% and outperforms Optimus by up to 47% in terms of Spearman R.

For the prediction of TE and EL, we used UTR-LM TE and UTR-LM EL and compared their performances with benchmark methods respectively^6–10^. Fig. 3d,e illustrates that UTR-LM TE and UTR-LM EL performs competitively with Cao-RF and outperforms other methods. Specifically, in terms of Spearman R, UTR-LM model outperforms Cao-RF by up to 5% and 8% for TE and EL tasks, and outperforms Optimus by up to 25% and 47% for TE and EL tasks. While Cao-RF proves effective in predicting TE and EL, it relies on more than 3000 handcrafted features including *k*-mer frequency, RNA folding energy, 5’ UTR length, and number of open reading frameworks, and its random forest model that may encounter scalability issues with larger datasets. In contrast, the training of a language model offers a more principled solution to modeling 5′ UTR sequences and only uses information (for example MFE) that can be easily computed. Thus, we believe that the UTR-LM provides a more robust and generalizable model for understanding 5’ UTR sequences.

### 2.3. UTR-LM identifies unannotated Internal Ribosome Entry Sites (IRESs) within the untranslated region

Internal ribosome entry sites (IRESs) are unique RNA sequences, most located within the 5’ UTR of mRNAs. Unlike the typical cap-dependent translation initiation that starts at the 5’ end of an mRNA, IRESs enable ribosomes to initiate translation directly at the internal sites. Approximately 10% of cellular and viral mRNA are believed to employ IRESs for translation initiation^22^. However, due to the limited number of verified IRESs, in-depth research and comprehension of IRES functionality have been hindered. In this study, we used the pre-trained language model for 5’ UTR sequences and transferred it to identify unannotated IRESs.

The goal of IRES prediction is to identify whether a given sequence can function as an IRES or not. For this purpose, we assembled a library of 46,774 sequences including both viral and cellular mRNAs, sourced from multiple databases^12, 23–26^ (see details in the Methods Section), with 37,602 sequences labeled as non-IRESs and 9,172 as IRESs. Building on the pre-trained UTR-LM, we developed a contrastive learning model to train a downstream IRES classifier, as elaborated in the Methods Section.

Several baseline methods exist for the task of predicting IRES, such as IRESfinder^13^, IRESpred^12^, IRESpy^11^, and DeepCIP^14^. IRESfinder is a logit model with framed *k*-mer features for cellular IRES prediction but may not be effective for viral IRES. DeepCIP is a specialized deep learning model designed specifically for circRNA IRES prediction (a subset of all IRES) and it uses both sequence and structure information. IRESPred is a support vector machine model for predicting both viral and cellular IRES based on 35 handcrafted features including sequence properties, structural properties, and interaction probabilities between 5’ UTR and small subunit ribosomal proteins. IRESpy is an XGBoost model that is trained on 340 global *k*-mer features, minimum fold energy, and 32 features including sequence properties and structural properties. For our comparison, we selected IRESpy as the benchmark because it represents the most recent method for both viral and cellular IRES detection and has shown advantages over IRESpred. We also compared our model with CNN baselines that were reported as top performers in previous literature^6, 7^. As illustrated in Fig. 4, the UTR-LM IRES classifier substantially outperforms the best-known benchmark, improving the AUPR from 0.37 to 0.52. Here we chose to focus on the metric AUPR because it is more suitable to evaluate classifiers on imbalanced data.

**Fig. 4.**
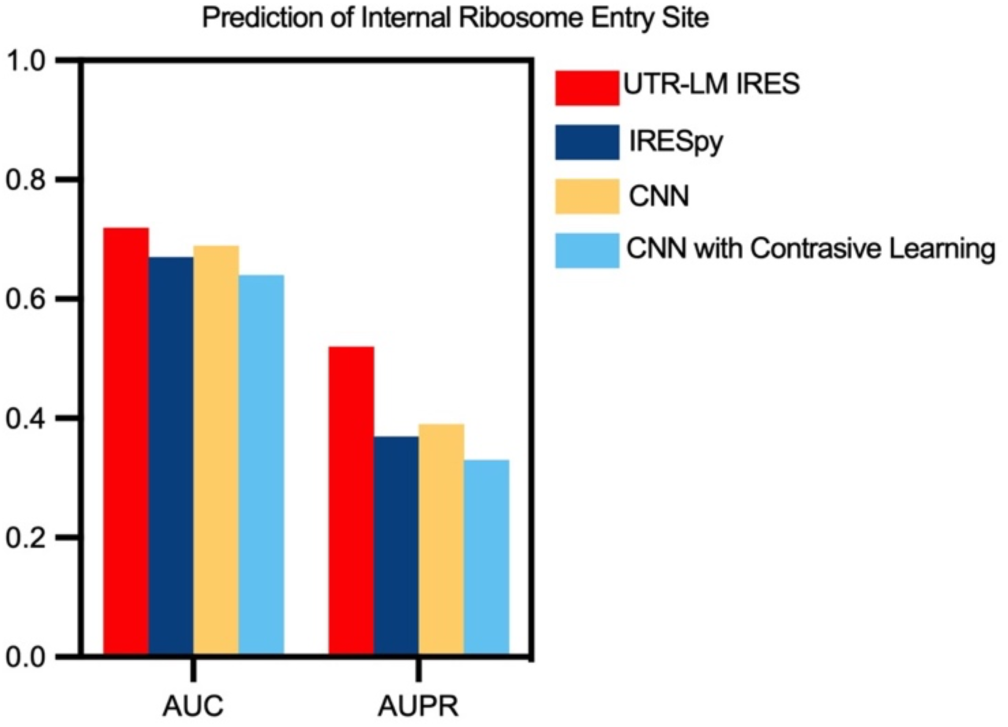
UTR-LM accurately identifies unannotated internal ribosome entry sites (IRES). Our method, the UTR-LM IRES classifier, achieves higher test accuracy than best-known baselines for this task. All models were trained and tested on a dataset of 46,774 sequences via 10-fold cross-validation.

### 2.4. Experimental assay finds novel 5’ UTR designs and confirms the validity of UTR-LM

Finally, we conducted a set of wet-lab experimental assay to validate our prediction model and generate novel 5’ UTR designs with high translation efficiency. We designed a library of 211 distinct 5’ UTR sequences with high predicted values of translation efficiency using the prediction model and independently from the human expert’s choice. In the experiment, we used expression of a luciferase reporter gene to measure mRNA translation in human cells (Fig. 5a). We first cloned all designed 5’ UTRs upstream of the same, standard luciferase reporter gene, and then transfected the synthesized mRNAs into C2C12 cells, followed by quantitative luciferase assay (Fig. 5b). Specifically, we measured the relative light units (RLU) that quantifies the luciferase activity. This allowed us to assess the protein production level from each mRNA, which provided a direct measurement of how the designed 5’ UTRs might affect the protein synthesis process. We also compared the predicted values of MRL and TE of the designed library with the background distribution (i.e., distribution of training data in each task). As visualized in Fig. 5c,d, the UTR-LM model predicts that these in-house designed 5’ UTRs exhibit significantly higher MRL and TE values than the background.

**Fig. 5.**
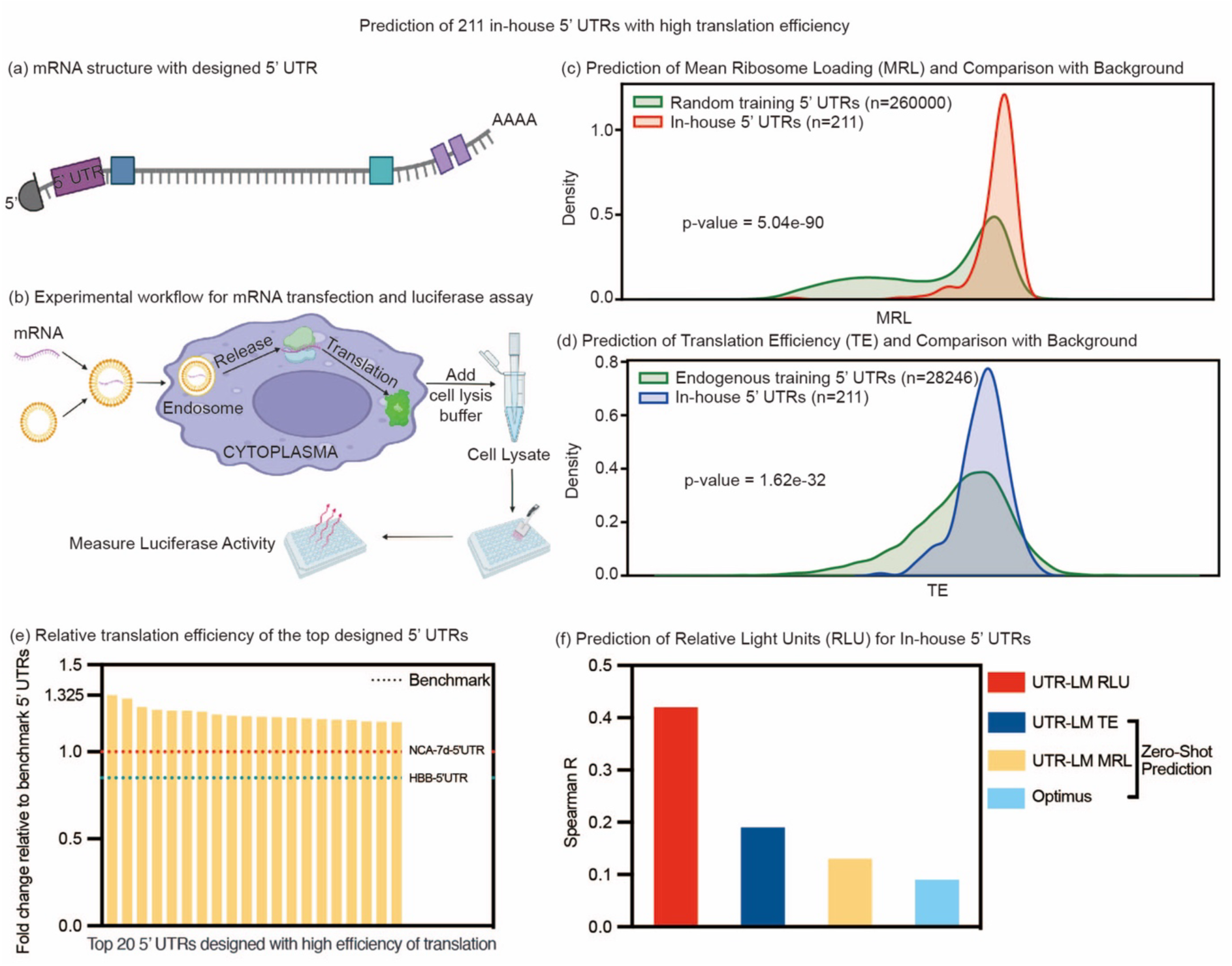
Experimental validation of UTR-LM model and top sequence designs generated by UTR-LM. A library of 211 in-house 5’ UTRs with high translation efficiency was designed and validated through wet-lab experiments. These sequences were subsequently used as an independent test set to verify the efficiency of our model. (a) The mRNA structure with in-house designed 5’ UTR. (b) Experimental workflow for mRNA transfection and luciferase assay. (c-d) The in-house 5’ UTRs had higher predicted values of mean ribosome loading (MRL) and translation efficiency (TE) compared to the background. (e) Wet-Lab Results: Relative translation efficiency of the top 20 designed 5’ UTRs compared to the benchmarks 5’ UTRs NCA-7d-5’UTR and HBB-5’UTR. (f) Using the wet-lab results as an independent test set, the UTR-LM gave substantially more accurate zero-shot predictions of the Relative Light Units (RLU) compared to the benchmark. Drawings in panels (a-b) were created with BioRender.com.

We next compared the efficiencies of the top in-house designed 5’ UTRs with well-established UTRs used for mRNA therapeutics. For benchmarks, we measured our designs against two well-known 5’ UTRs, namely the HBB-5’UTR^27^ and the NCA-7d-5’UTR^15^. The HBB-5’UTR, a 5’ UTR from the human hemoglobin subunit beta, is commonly used in studies of mRNA translation and stability^27^. The NCA-7d-5’UTR, an optimized 5’ UTR for protein-coding mRNAs, has demonstrated effective delivery via lipid-derived TT3 nanoparticles, resulting in pronounced expression of potential SARS-CoV-2 antigens^15^. Our experiment validated that top candidates in our designed library achieved significantly improved protein production level (Fig. 5e). In particular, the top 5’ UTR sequence found in our assay had a 32.5% increase in protein production level compared to NCA-7d-5’UTR. These results further validated our prediction model, showing its potential to generate novel designs for mRNA vaccines and therapeutics. Full details of the experimental design and methods are given in Supplementary Materials A.5.

Next, we evaluated our UTR-LM for zero-shot fitness prediction using our in-house design library and wet-lab results as an independent test set. A challenge is that our experiment did not give direct measurements of TE or MRL. Thus, we used the RLU as the prediction target, measured by the fold change of log2-transformed RLU relative to NCA-7d-5’UTR as the ground-truth label (referred to as NCA-7d-5’UTR_FC_log2RLU). For zero-shot fitness prediction, we transferred the learnt models UTR-LM MRL and UTR-LM TE to apply to the new dataset, without any additional training or fine-tuning, and tested their performances on the new unseen labels. We also compared our results with the benchmark model, Optimus^7^, which was trained on MRL. We note that the training labels MRL and TE used in our model were different from the new prediction target RLU, but they are believed to be highly correlated. To evaluate the performance, we computed the Spearman rank coefficient R (i.e., Spearman R). As illustrated in Fig. 5(f), UTR-LM MRL and UTR-LM TE outperformed the benchmark Optimus, with UTR-LM TE exceeding Optimus by more than one fold, demonstrating the transferability of our model across tasks and modalities.

Further, we applied the pre-trained UTR-LM embedding, trained a new RLU predictor using our in-house data and tested it via 10-fold cross-validation. As illustrated in Fig. 5f, our UTR-LM RLU model substantially outperformed the benchmark Optimus and the UTR-LM without retraining. The result emphasizes the considerable potential of the language model approach in modeling genome sequences, demonstrating its robustness and generalizability.

Lastly, we sought to analyze if the UTR-LM model increases our chance of identifying the top 5’ UTR designs. For this purpose, we first picked an arbitrary quantile level, say top x%. Then we compared the top x% designs in our in-house library ranked by predicted value with the top x% designs ranked by wet-lab measurement. Let y denote the percentage of overlap between the two sets. Here, we focus on the ratio y/x to quantify the “success factor” for finding top x% designs using our prediction model. For example, if the prediction model is purely random, the expected success chance factor would equal to 1.0. In Extended Fig. 1, we visualized the success factors computed using comparable models, including UTR-LM MRL, UTR-LM TE, and Optimus, across different values of x. It shows that our pre-trained language model significantly improves the success rate of finding top 1%-15% designs by 2-4 folds.

## 3. Discussion

### 3.1. The pre-trained embedding of UTR-LM recognizes underlying patterns

We examine the 5’ UTRs across the five species used for pretraining. In Fig. 6a, we illustrated the frequency of each nucleotide at each base, per specie. One can see that rat, mouse and humans are highly similar, while chicken and zebrafish are substantially different from the former ones. We computed the silhouette scores, a metric that measures the separation between clusters, for pairs of species using both the UTR-LM embedding and the 4-mer representation. As shown in Fig. 6b, our UTR-LM embeddings achieved significantly higher silhouette scores than those of the 4-mer representation. The results suggest that our pretrained language model provides more meaningful representations for differentiating between species. Additionally, we visualized the pre-trained embeddings from the five species and examined their minimum fold energy (MFE) values. As shown by Fig. 6c, the UTR-LM embedding of 5’ UTR captured most variation in the MFE value, while the traditional 4-mer representation struggled.

**Fig. 6.**
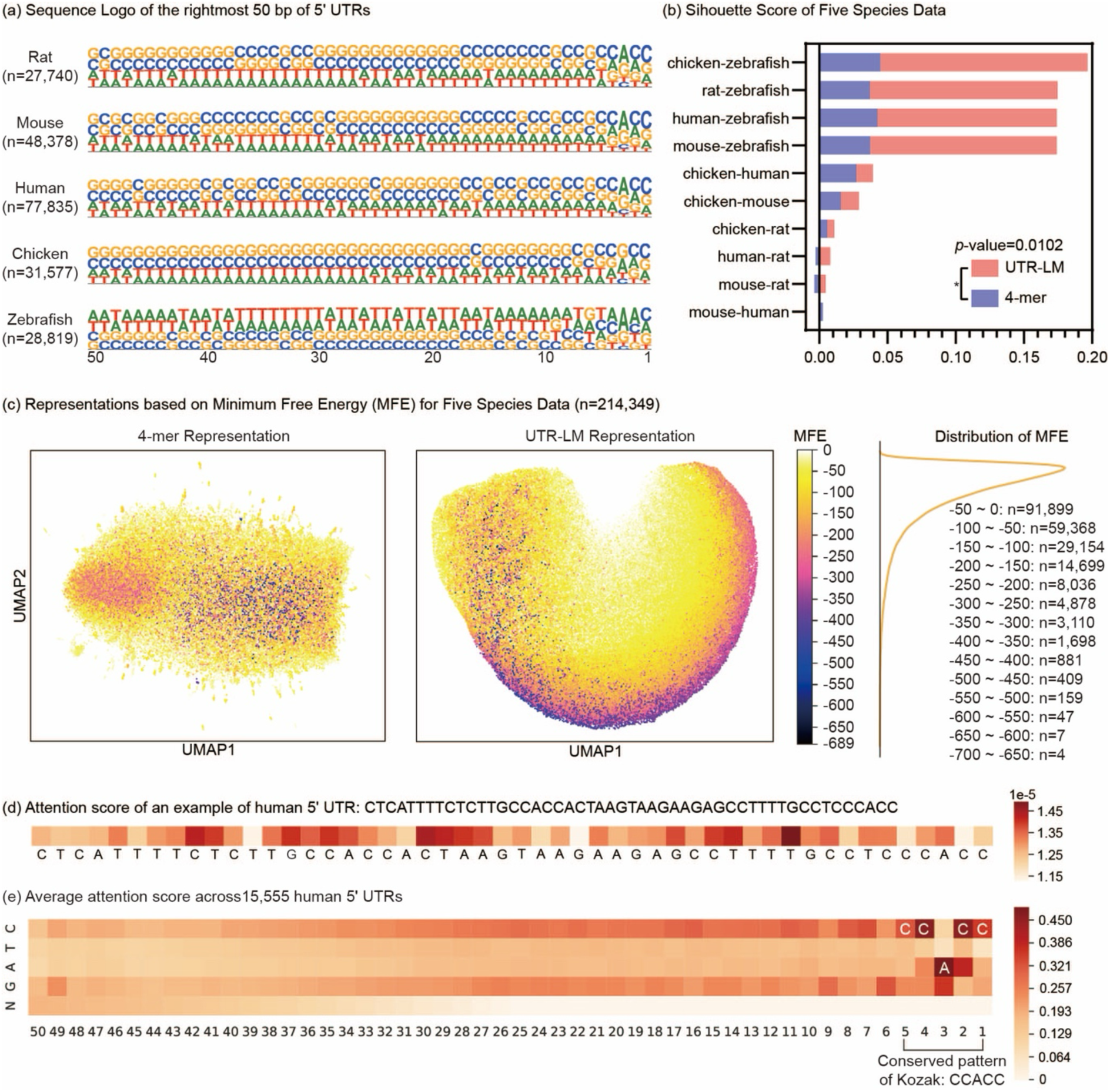
The UTR language model recognizes underlying patterns and reveals known motif patterns. (a-c) The pre-trained UTR-LM embedding outperforms the traditional 4-mer representation in differentiating between species and captures the minimum free energy (MFE) patterns. Analyses were based on 5’ UTRs from five species sourced from the Ensembl database. (a) Sequence logo representing the last 100-bp of the pre-trained 5’ UTRs for each of the five species. (b) Silhouette scores computed using the embedding indicates the language model is better than 4-mer representation for differentiating species. (c) Visualization of 4-mer representation and UTR-LM embedding, colored by the MFE. (d) Visualization of sequence-level attention of a human 5’ UTR example, highlighting the significance of each site in the sequence to its mean ribosome loading. (e) Visualization of position-level patterns within the 15,555 human 5’ UTRs, where potential motifs are specifically highlighted based on attention. The Kozak consensus sequence, which is typically accepted as the conserved pattern CCACC, has been recognized as a motif that largely affects the ribosome loading. The x-axis represents the position inside the 5’ UTRs.

### 3.2. The attention-based motif detection uncovers proven patterns

The attention mechanism in the transformer architecture helps to draw connections between any parts of the sequence. A high attention score for a specific site suggests it could be influential in determining the target function. For example in Fig. 6d, we visualized the attention scores of a human 5’ UTR associated with the MRL prediction.

Next, we examined the attention scores of UTR-LM with the hope of identifying potential patterns of 5’ UTRs. We refer to the Methods Section for computation details. We analyzed the average attention score per each position and per nucleotide cross all human 5’ UTRs, as shown in Fig. 6e. Those high attention scores in positions 1-6 revealed the Kozak consensus sequence (KCS)^16, 17^, which is a nucleotide motif that functions as the protein translation initiation site in most mRNA transcripts. Within the KCS region, the attention scores are align well with the proven conserved nucleotide patterns^28^. Specifically, the attention score for each nucleotide at each position is consistent with their proven frequencies in the literature^28^. Moreover, our analysis detected the typically accepted conserved pattern CCACC^29^ (positions 1-5 in Fig. 6e) among KCS variations, and recognized the fact that nucleotides G and C are generally more impactful than A and T within the KCS^16^.

### 3.3. Summary

We introduced a language model for 5’ UTRs that integrates sequence, secondary structure, and MFE via semi-supervised training. The UTR-LM model has learned meaningful semantic representations from 5’ UTR sequences that pertain to the mRNA translation process. In particular, it applies to predicting the mean ribosome loading (MRL), translation efficiency (TE), mRNA expression levels (EL), and internal ribosome entry site (IRES) and outperforms the best-known baseline in each task.

In summary, our method provides a novel and effective approach for modeling the 5’ UTR and is well-suited for studying the downstream translation process. Additionally, we generated 211 in-house 5’ UTR designs and measured their performances via wet-lab experiments. Our experiment revealed a set of highly-efficient novel 5’UTR designs with potential therapeutic value and further validated the prediction model.

## 4. Methods

### 4.1. Overview of UTR-LM

We developed a unified foundation language model to provide meaningful and rich representations for 5’ UTRs. The model adopts a transformer architecture and it is trained using multi-modal data, including raw sequences, secondary structure (SS), and minimum free energy (MFE). The model is pre-trained in a semi-supervised learning manner via mask reconstruction, SS prediction and MFE prediction. It is later fine-tuned for a variety of downstream function prediction tasks, where it is shown to improve the state-of-the-art performances in each task.

We trained UTR-LM on two computing clusters. The first cluster, hosted on the Amazon Web Services cloud platform, was equipped with four Tesla V100-SXM2 GPUs, each boasting 16 GB of high-bandwidth memory. The second cluster, the Stanford University’s Sherlock high-performance computing system, employed four TESLA_P100_PCIE GPUs, each with 32 GB of memory. We configured the training to a maximum of 200 epochs and two days.

### 4.2. Datasets

For pre-training the language model, we collected unlabeled 5’ UTR sequences from three sources: the Ensembl database^30^, synthetic libraries from Sample et al.^7^, and endogenous human 5’ UTR data analyzed by Cao et al.^8^ We pre-processed the raw data to keep only high-quality and well-defined 5’ UTRs, details are shown in Supplementary Materials A.1. After the pre-processing steps, the final large-scale dataset from several sources is obtained:

We obtained 214,349 unlabeled 5’ UTR sequences from the Ensembl database^30^, spanning five species: human, rat, mouse, chicken, and zebrafish.

We obtained eight synthetic libraries of random 5’ UTRs truncated to 50 nucleotides long from Sample et al.^7^ These libraries are sorted into two groups based on their coding sequence (CDS): six libraries are linked to the enhanced green fluorescent protein (eGFP), and two are linked to mCherry. The eGFP-linked libraries consist of two unmodified uridine (U) libraries (U_1_ and U_2_), two pseudouridine (Ψ) libraries (Ψ_1_ and Ψ_2_), and two 1-methyl pseudouridine (m_1_Ψ) libraries (m_1_Ψ_1_ and m_1_Ψ_2_). Each eGFP library contains approximately 280,000 distinct 5’ UTRs. The mCherry-linked libraries, named mC-U_1_ and mC-U_2_, each contain around 200,000 unique 5’ UTRs.

We obtained three endogenous human 5’ UTR datasets analyzed by Cao et al.^8^, each originating from a distinct cell line or tissue type: human embryonic kidney 293T (HEK), human prostate cancer cell (PC3), and human muscle tissue (Muscle). The HEK, PC3, and Muscle datasets comprised 14,410, 12,579, and 1,257 sequences, respectively.

In addition, we also included unlabeled raw sequences from datasets of downstream tasks in the pretraining. For more in-depth description of each dataset, please see Supplementary Materials A.

### 4.3. Architecture and pre-training of UTR-LM

We developed a specialized language model for studying 5’ UTRs, called UTR-LM (as illustrated in Fig. 1). The main architecture comprises an encoder block and a predictor block. The encoder block consists of a six-layer transformer^31^ with 16 multi-head self-attention. The layer normalization and residual connections are applied before and after each encoder block. The predictor block is a two-layer feed-forward neural network.

In our UTR-LM model, a 5’ UTR sequence of length L is input as a series of nucleotide tokens (such as ‘A’, ‘G’, ‘C’, ‘T’), along with a special [CLS] token. These tokens are first converted into 128-dimensional vectors via an embedding layer, forming an (L+1) × 128 matrix. This matrix then goes through the encoder block, generating a representation for each nucleotide token in the sequence.

In the pre-training stage, we used a mix of self-supervised learning and supervised learning. For the self-supervised training part, we followed the masked language modeling (MLM)^32^, where we randomly masked 15% of the nucleotide tokens in the 5’ UTR sequence. The model is then trained to predict these masked tokens by minimizing the cross-entropy loss to encourage accurate reconstruction. We refer to this part as the masked nucleotide (MN) training task. This training objective function is

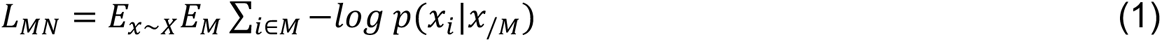

In equation (1), we randomly select a set of indices, denoted as 𝑀, from each input sequence 𝑥 (making up 15% of the entire sequence). The token at each index 𝑖 is replaced with the mask token [MASK]. The objective is to minimize the negative log-likelihood of the correct reconstruction of each 𝑥*_i_*, when the unmasked part 𝑥_/M_ is given as context.

For the supervised training part, we used two labels to provide auxiliary supervision. First, we included the SS of 5’ UTR for training, which is calculated using the software ViennaRNA^33^. We represented the SS using the "dot-bracket" notation^33^, where paired nucleotides are given by "(" and ")" characters and unpaired nucleotides are given by a ".". For instance, the SS of the sequence AUGCAUGCGAUCAGC is given by "(((..)))..((.))". To utilize the SS, we introduced an MLM-inspired task (which we call the SS task) to predict the SS symbols associated with masked nucleotides. The training objective function is

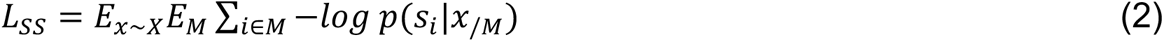

equation (2) differs from equation (1) in that it uses the SS 𝑠 as input. It attempts to reconstruct the masked SS, when the unmasked part 𝑥_/M_ is given as context. The set of mask indices 𝑀 is the same as for the MN task. Second, we used the MFE of the 5’ UTR as an additional target of prediction, because there exists a proven correlation between the MFE and the translation efficiency^7^. We refer to this step as the MFE training task, where the predictor block uses the [CLS] token’s representation to estimate the MFE value. We used the ViennaRNA software^33^ to calculate the actual MFE. The training objective is to minimize the mean squared error:

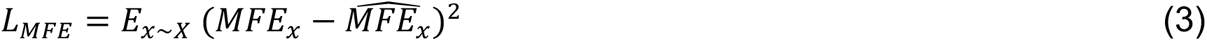

where 𝑀𝐹𝐸*_x_* and 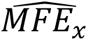 are the actual and predicted MFE of the input sequence 𝑥.

### 4.4. Downstream training strategies

We used the learned representations from UTR-LM for several downstream tasks, as illustrated in Fig. 1b,c. We extended the predictor block of the pre-trained UTR-LM from predicting MFE to predicting the downstream label. Instead of using UTR-LM merely as a static generator of continuous embeddings from discrete nucleotide sequences, we used the pre-trained parameters as an initial point. Then, we fine-tuned the entire model, adjusting all its parameters.

#### 4.4.1. Prediction of Mean Ribosome Loading (MRL)

Ribosome loading, a measure of mRNA translation efficiency influenced by factors like the 5’ UTR, is experimentally assessed through techniques like polysome profiling to derive the mean ribosome loading (MRL). Understanding and designing the 5’ UTR can optimize MRL, thereby enhancing protein production for applications such as biotechnology and therapeutics. Here, we attempted to predict MRL based on the 5’ UTR sequence.

Firstly, we used eight libraries of randomized 50-nucleotide oligomer 5’ UTRs^7^, which are U_1_, U_2_, Ψ_1_, Ψ_2_, m_1_Ψ_1_, m_1_Ψ_2_, mC-U_1_ and mC-U_2_. These libraries provide a measure of MRL per sequence. Details and statistics can be found in Supplementary Material A.3. Followed by Sample et al.^7^, for each library, we established specialized version of UTR-LM model, which we collectively refer to as UTR-LM MRL. Each version was initially pre-trained on both the Ensembl database and its respective library. We then fine-tuned these specialized models, updating all parameters for better performance.

We performed independent tests on a dataset containing 83,919 random 5’ UTRs and another one containing 15,555 human 5’ UTRs, both with varying lengths from 25 to 100 bp. We employed two testing strategies for independent tests. The first strategy, proposed by Optimus^7^, employed a length-based held-out testing approach. Each test set contains 7,600 random 5’ UTRs or 7,600 human 5’ UTRs with the most reads. The remaining 76,319 random 5’ UTRs were used as the training set for both independent tests. The second strategy is to use a 10-fold cross-validation. Because the U_1_ is commonly used in prior UTR research^6, 7^, we started with the UTR-LM MRL model that was trained on U_1_ as the initial model and then finetuned it using the training set.

#### 4.4.2. Prediction of mRNA Expression Level (EL) and Translation Efficiency (TE)

Protein production is influenced by mRNA expression level (EL), measured in RNA-seq RPKM^8^, and translation efficiency (TE), calculated as the ratio of Ribo-seq to RNA-seq RPKM^8^. Both factors are essential for understanding how the 5’ UTR influences the rate at which mRNA is translated into protein.

We used three different human datasets^8^ for the TE and the EL tasks. These datasets are named Muscle, PC3, and HEK, and together they contain 41,446 unique 5’ UTRs. Each sequence of these datasets provides measurements of TE and EL. In alignment with the limitation of commercially available single-stranded DNA template biosynthesis^8^, a fixed 5′ UTR length of 100-bp was chosen for training. More detailed statistics can be found in Supplementary Materials A.4.

For each dataset, we trained a TE prediction model and EL prediction model, respectively. The model is fine-tuned based on the parameter of the UTR-LM model pre-trained on both Ensembl database and 41,446 human 5’ UTRs as an initial point.

#### 4.4.3. Identification of Internal Ribosome Entry Site (IRES) with Contrastive Learning

Most internal ribosome entry sites (IRESs) are unique sequences found within the 5’ UTRs of mRNAs. These specialized sequences enable cap-independent translation initiation and are thought to regulate translation for a subset of cellular and viral mRNAs^22^.

We assembled an unbalanced dataset sourced from several studies, including Weingarten-Gabbay et al.^23^, IRESbase^24^, IRESite^25^, Rfam^26^, and IRESpred^12^. This assembled dataset contains 46,774 sequences, with 37,602 sequences labeled as non-IRESs and 9,172 identified as IRESs. More detailed statistics can be found in Supplementary Materials A.5.

We adopted a contrastive learning approach to distinguish between IRES and non-IRES categories. Firstly, we constructed contrastive samples where each sample consists of a pair of a IRES and a non-IRES sequence. For each sequence in the training set, we made one contrastive sample by randomly selecting a sequence with the opposite label to pair with it.

Although many IRES are often found in the 5’ UTR of mRNA, they can also appear elsewhere. To address this, we transferred the UTR-LM model to focus specifically on IRES. We started by using the U_1_-trained version of UTR-LM as our initial base model. Then, we fine-tuned this model by minimizing the combination of three loss functions. The first is the cross-entropy loss of the MN task, illustrated as equation (1). The second is a binary cross-entropy loss for IRES vs. non-IRES classification, formulated as follows:

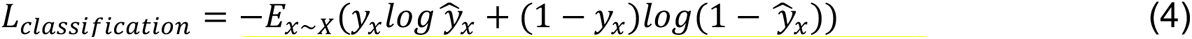

where 𝑦*_x_* and 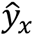 stand for the ground-truth label and the predicted probability of the label being IRES for the sequence 𝑥, respectively. The third is a specially designed contrastive loss that measures the difference between the predicted probabilities for sequences labeled as non-IRES and sequences labeled as IRES. Therefore, the loss function is computed over contrast samples, which contain pairs of an IRES and a non-IRES. It is formulated as follows:

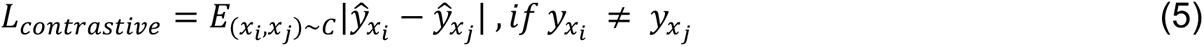

where the pair (𝑥*_i_*, 𝑥_j_) consists of two sequences selected from the set of contrastive samples, denoted by 𝐶. Since the pair comes from contrastive samples, their ground-truth labels are not equal 𝑦*_x_*_!_ ≠ 𝑦*_xj_*. The variables 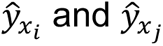 represent the predicted probabilities of the label being IRES for sequences 𝑥*_i_* and 𝑥_j_, respectively.

### 4.5. Attention-based motif detection

Attention scores provide insights into the contribution of specific positions in the input sequence to the model’s prediction. Utilizing the attention scores from UTR-LM allows us to pinpoint crucial sequence motifs in 5’ UTRs. At the sequence-level, we examined the attention scores across individual sequences to find regions that highly contribute to the predictive result. At the position-level, we averaged attention scores from the same nucleotide positions across multiple sequences, revealing potential patterns that contribute similarly in different 5’ UTRs. The computation workflow is given in Extended Fig. 2.

To capture attention information at the sequence level for an individual 5’ UTR, we adopted a methodology inspired by ESM^34^. We generate an initial attention matrix for each sequence, which has dimensions (# layers × # heads, L, L), where # layers and # heads are hyper-parameters of the self-attention layer, and L is the sequence length. We first make the attention matrix symmetrical by summing it with its transpose. We then normalize the symmetrized matrix across the (L, L) dimension using the formula:

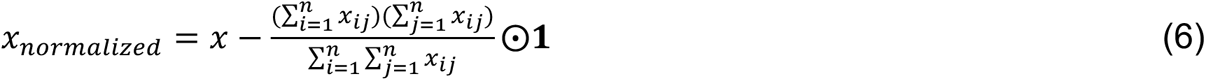

equation (6) describes how to perform element-wise subtraction to compute a normalized matrix 𝑥*_normalized_* from an input matrix 𝑥. Here, ⊙ denotes element-wise multiplication, and 𝟏 is a matrix of ones with dimensions matching 𝑥. This allows the average value to be broadcasted across all elements of 𝑥 during subtraction. Specifically, the average value is calculated by taking the product of the sums of the rows 𝑖 and columns 𝑗 of 𝑥 and dividing it by the sum of all elements in 𝑥. Next, we sum the normalized matrix across the # layers × # heads dimension, thereby reducing the dimensions to (L, L). Finally, we sum along the rows of the resulting (L, L) matrix to generate the final attention vector for the 5’ UTR sequence.

To analyze attention at the positional level, we generated attention vectors for 15,555 human sequences. Across these different sequences, we computed the average attention score for each nucleotide type at each specific position. This provides insight into how specific nucleotides at particular positions contribute to the overall sequence behavior.

### Data Availability

The datasets are available and can be downloaded at https://drive.google.com/drive/folders/1oGGgQ33cbx340vXsH_Ds_Py6Ad0TslLD?usp=sharing. This link includes training data for the pre-trained model as well as datasets for various downstream tasks. Detailed statistics for these datasets are provided in Supplementary Materials A.

### Code Availability

The code is freely available at https://github.com/a96123155/UTR-LM under the GNU General Public Licence Version 3. This repository includes dependencies, the operating environment, usage instructions, and interactions between the code and its results. The corresponding DOI for the project is https://doi.org/10.1234/utrlm.

### Webserver Availability

The webserver is freely available at https://huggingface.co/spaces/Shawn37/UTR_LM.

## Acknowledgements

We would like to extend our heartfelt thanks to Stanford University, Princeton University, and RVAC Medicines for their invaluable support throughout this research.

---

**Extended Fig. 1.**
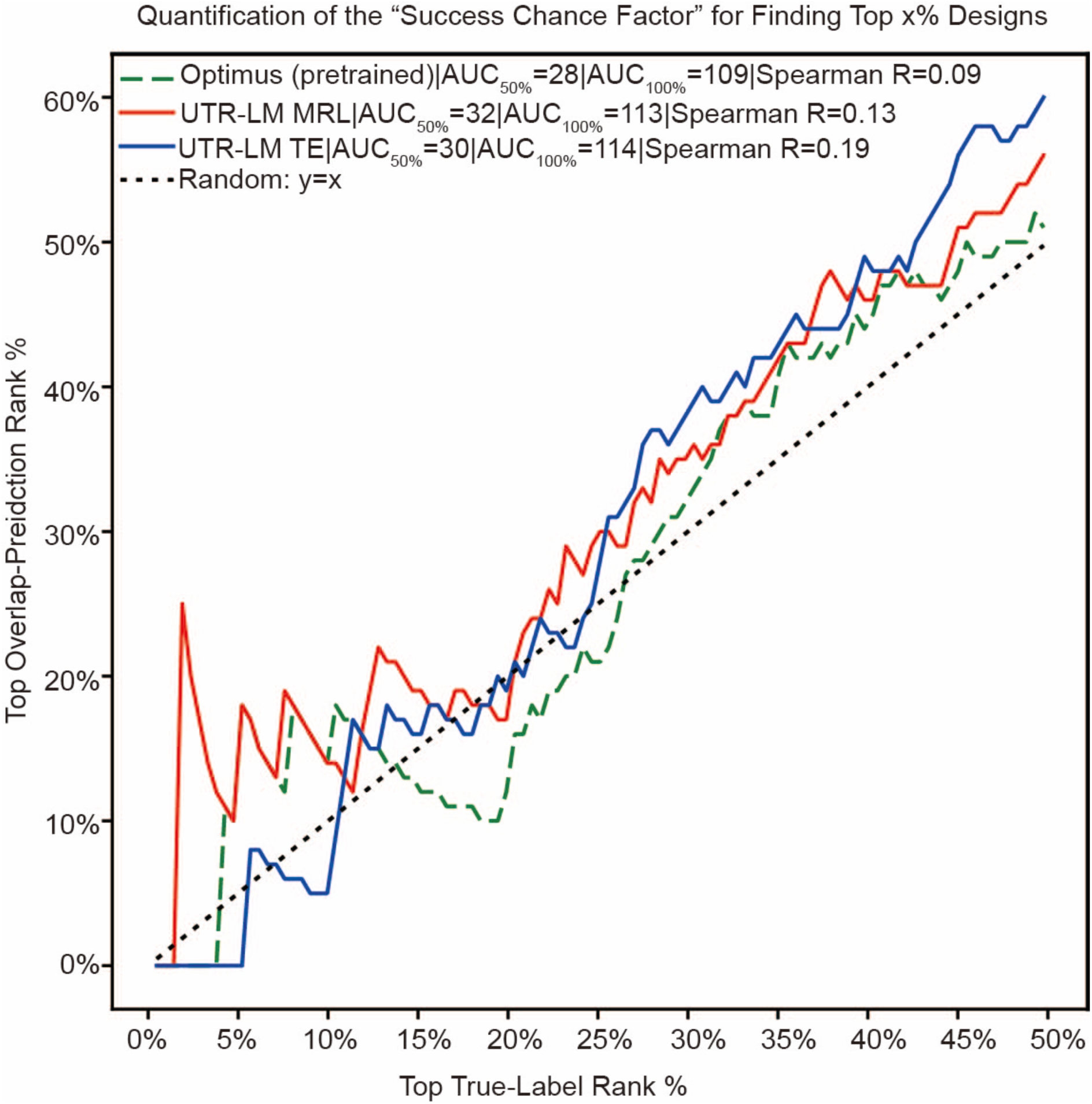
Prediction of NCA-7d-5’UTR_FC_log2RLU for the designed 5’ UTRs using UTR-LM and Optimus without retraining. This serves to quantify the “success chance factor” when identifying the top x% of designs. The x-axis denotes the arbitrary quantile level (i.e., top-x%) based on the ground-truth label. We compare the top x% designs in our in-house library ranked by predicted value with the top x% designs ranked by wet-lab measurement. The y-axis denotes the percentage of overlap between the two sets.

**Extended Fig. 2.**
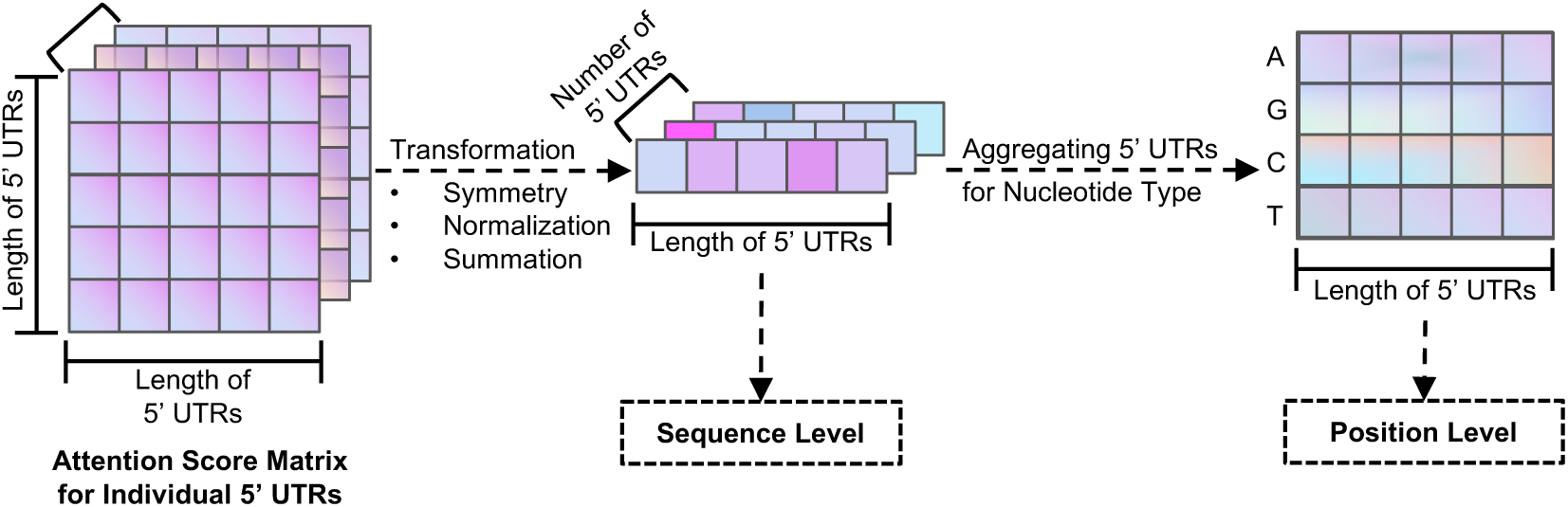
The computation flow of identifying patterns in 5’ UTR sequences based on attention scores.

## Supplementary Materials

## A. Data Description

### A.1. Pre-processing the data

Initially, we removed any coding sequence (CDS) or non-5’ UTR fragments from the raw sequences. Next, we identified and removed duplicate sequences. Then, we truncated the sequences to fit within a range of 30 to 1022 bp, following the guidelines from previous work established by Cao et al.^8^ and ESM^34^. Lastly, we filtered out incorrect or low-quality sequences following the work established by Sample et al.^7^ and Cao et al.^8^

### A.2. Data of five species from the Ensembl database

We collected 214,349 unlabeled 5’ UTR sequences from the Ensembl database^30^, spanning five species: human, rat, mouse, chicken, and zebrafish. These species have high-quality and manual gene annotations. The potential benefits of incorporating 5’ UTRs from an additional 309 species, which have comparatively lower-quality gene annotations, remain an avenue for future exploration. Detailed statistics for the data from these five species can be found in Supplementary Table 1. After data cleaning, the counts of 5’ UTRs from the Ensembl database^30^ are 77,835, 27,740, 48,378, 31,577, and 28,819, respectively. We chose these species as they represent a diverse spectrum of vertebrates with varying evolutionary connections. The details can be found in Supplementary Table 1.

**Supplementary Table 1.** Endogenous data collected from the Ensembl database. The min and max of sequence lengths are 30 and 1022, respectively.

| Species | Counts (#) | # With ATG | # Without ATG | Mean of Length | Median of Length |
| --- | --- | --- | --- | --- | --- |
| Human | 77,835 | 48,128 | 29,707 | 249 | 190 |
| Rat | 27,740 | 17,639 | 10,101 | 309 | 201 |
| Mouse | 48,378 | 29,624 | 18,754 | 234 | 178 |
| Chicken | 31,577 | 20,999 | 10,578 | 233 | 193 |
| Zebrafish | 28,819 | 21,816 | 70,03 | 240 | 172 |
| All | 21,4349 | 13,8206 | 76,143 | 250 | 186 |

### A.3. Randomized 50-nucleotide oligomer data with mean ribosome loading

Sample et al.7 proposed eight distinct 5’ UTR libraries, each containing random 50 nucleotide sequences, to evaluate translation rules using mean ribosome loading (MRL) measurements. These libraries are called as U_1_, U_2_, Ψ_1_, Ψ_2_, m_1_Ψ_1_, m_1_Ψ_2_, mCU_1_, and mC-U_2_. In this context, the U, Ψ, m_1_Ψ, mC symbolize unmodified uridine, pseudouridine, and 1-methyl pseudouridine, and the CDS of the mCherry, respectively.

Specifically, the mC-U1 and mC-U2 libraries, containing about 200,000 5’ UTRs each, are based on the mCherry CDS. The remaining libraries utilize the CDS of the enhanced green fluorescent protein (eGFP) and each comprises roughly 280,000 5’ UTRs. Further details are provided in Supplementary Table 2.

Sample et al.7 generated 83,919 random 5’ UTR sequences and 15,555 human 5’ UTRs, covering a length range from 25 to 100 nucleotides. The number of total reads per 5’ UTR in both libraries is greater than 10.

**Supplementary Table 2.** Eight randomized 50-nucleotide oligomer data with mean ribosome loading.

| Library | CDS | Uridine | Counts (#) | # With ATG | # Without ATG |
| --- | --- | --- | --- | --- | --- |
| $U_1$ | eGFP | Unmodified | 280,000 | 151,059 | 128,941 |
| $U_2$ | eGFP | Unmodified | 280,000 | 150,114 | 129,886 |
| $mC-U_1$ | mCherry | Unmodified | 280,000 | 104,643 | 95,357 |
| $mC-U_2$ | mCherry | Unmodified | 280,000 | 103,975 | 96,025 |
| $\Psi_1$ | eGFP | Pseudouridine | 279,968 | 149,370 | 130,598 |
| $\Psi_2$ | eGFP | Pseudouridine | 280,000 | 149,648 | 130,352 |
| $m_1\Psi_1$ | eGFP | 1-methylpseudouridine | 279,968 | 149,246 | 130,722 |
| $m_1\Psi_2$ | eGFP | 1-methylpseudouridine | 280,000 | 147,908 | 132,092 |

### A.4. Human 5’ UTR data for translation efficiency

We adopted the endogenous human 5’ UTR data analyzed by Cao et al.^8^, which involves matching public RNA-Seq and Ribo-Seq datasets. Data from three distinct cell lines/tissues, specifically human embryonic kidney 293T (HEK), human prostate cancer cell (PC3), and human muscle tissue (Muscle), were assembled into three separate endogenous datasets. The mRNA expression level (EL) was determined using RNA-seq RPKM (Reads Per Kilobase of transcript per Million mapped reads), and the mRNA translation efficiency (TE) for each transcript was calculated by dividing the Ribo-seq RPKM by the RNA-seq RPKM^8^. In alignment with the limitation of commercially available single-stranded DNA template biosynthesis^8^, a fixed 5’ UTR length of 100-bp was chosen for algorithm training. Transcripts that fail to meet the minimum coverage criteria are excluded, specifically those with RNA-Seq RPKM < 5 and Ribo-Seq RPKM < 0.1 for the gene. The HEK, PC3, and Muscle datasets comprised 14,410, 12,579, and 1,257 sequences respectively. For each 5’ UTR, we adopted the 100-bp sequence upstream of the CDS, covering 29% of human 5’ UTRs. More detailed statistics can be found in Supplementary Table 3.

**Supplementary Table 3.** Three datasets of human endogenous 5’ UTRs with mRNA translation efficiency and expression level.

| Source | Number (#) | # With ATG | # Without ATG | Mean of Length | Median of Length |
| --- | --- | --- | --- | --- | --- |
| Muscle | 1257 | 613 | 644 | 183 | 126 |
| PC3 | 12,579 | 6,075 | 6,504 | 192 | 132 |
| HEK | 14,410 | 7,018 | 7,392 | 185 | 127 |

### A.5. Dataset for the Internal Ribosome Entrance Site

We assembled a dataset of sequences, each labeled binary based on their function as the internal ribosome entry site (IRES). This dataset was sourced from several studies, including those by Weingarten-Gabbay et al.^23^, IRESbase^24^, IRESite^25^, Rfam^26^, and the training dataset from IRESpred^12^. To maintain data uniqueness, we removed entries that were redundant across the different sources. The finalized dataset contained 46,774 entries. It is important to highlight that this dataset encompasses sequences with diverse lengths, from 6 to 1,730 nucleotides. Additionally, the label distribution is extremely unbalanced, with 37,602 sequences labeled as non-IRESs and 9,172 identified as IRESs.

### A.6. In-house UTR design and experimental validation

#### A.6.1. 5’ UTR sequence design by genetic algorithm

All sequence evolutions started with randomized sequences. Over the course of 100 iterations, either a single base or two bases (with a 50% probability) were randomly chosen in each iteration and mutated. The fitness of the sequence was then evaluated. If the new sequence had a performance that was higher or closer to the target, it was retained; otherwise, the original sequence was kept. The fitness was assessed by Optimus re-trained on MRL and TE tasks. From multiple rounds of sequence evolution, 211 sequences with high predicted MRL and TE values were selected.

#### A.6.2. Plasmid synthesis and in vitro transcription

The designed 5’ UTRs were synthesized and inserted into a custom plasmid backbone, positioned before the codon-optimized coding sequence of the luciferase gene and after the T7 promoter. A DNA template for in vitro transcription (IVT) was produced via plasmid linearization using BspQI restriction enzyme. For the IVT reaction, we utilized the HiScribe T7 high-yield RNA synthesis kit (NEB). As a cap structure analog, we employed the 3′-O-Me-m7 G(5′) ppp(5′)AmG RNA cap (TriLink Biotech). And the DNA template was digested with DNase I (NEB), and the IVT mRNA was purified using the Monarch® RNA Cleanup Kit (NEB).

#### A.6.3. Cell culture

C2C12 mouse myoblasts (ATCC, CRL-1772) were cultivated in DMEM with 10% FCS and 1 mM sodium pyruvate. HepG2 human hepatocellular carcinoma (ATCC, HB-8065) were cultivated in DMEM with 10% FBS and 1% Pen-Strep. All cells were cultivated at 37°C and a CO2 content of 5%.

#### A.6.4. mRNA transfection and luciferase Assay

Prior to day of transfection, C2C12 and HepG2, 20000 cells/well were cultured with 100 μl of complete DMEM growth medium seed in 96-well tissue culture plate. In the following day, cells with density about 50∼80% confluent were transfected with 25, 12.5, 6.25, 3.125ng of mRNA per well using Lipofectamine™ MessengerMAX™ Transfection Reagent (Invitrogen) according to manufacturer’s protocol. Then, cells were incubated at 37°C in a CO2 incubator. In vitro luciferase expression was measured at 16h and 48h post transfection using the One-Glo Luciferase Assay System (Promega). The efficacy of translation of different mRNA constructs was calculated as fold change relative to the benchmark mRNA, NCA-7d-5’UTR^15^, which comes from a pair of optimized 5’ and 3’ UTRs. To simplify, we only used the results from the 6.25ng dose at 16h in the C2C12 cell line to evaluate the machine learning methods.

## B. Details of Training

### B.1. Hyper-parameters of pre-training process

**Supplementary Table 4.** Data for pre-training process. The pre-training tasks labeled as MN, SS, and MFE correspond to the randomly 15% masked token, secondary structure, and minimum free energy tasks, respectively. Meanwhile, the downstream tasks identified as MRL, TE, and EL refer to mean ribosome loading, translation efficiency, and mRNA expression level tasks, respectively.

| Data Source | MN task | SS task | MFE task |  |  |
| --- | --- | --- | --- | --- | --- |
|  |  |  | MRL task | TE task | EL task |
| Ensembl Database | √ | √ |  |  |  |
| Randomized 50-nucleotides oligomer library | √ | √ | √ |  |  |
| Human datasets from tissue/cell lines | √ | √ |  | √ | √ |

### B.2. Three types of representations generated by pre-training model

The per-token representation consists of a list of vectors, with each vector corresponding to a single token in the input sequence. This representation captures details about the token, such as its part-of-speech, syntactic dependencies, and other relevant features. Because these vectors are derived from applying an attention mechanism to the hidden states, they highlight the importance of each token for the final prediction.

The mean representation is a single vector derived by averaging all the per-token representations within a sentence, serving as a summary representation of the input sequence.

A distinct token, termed the [CLS] token, is positioned at the beginning of the sequence. Its representation serves as a learned vector that encapsulates the entire input sequence and is suitable for sequence classification or regression. The proposed pre-training model integrates supervised information as auxiliary labels, aiming to derive a representation tailored to specific tasks. This representation not only capture the global information of the input sequence, but also extract the hidden information of specific downstream tasks. Employing this representation proves beneficial in generating accurate predictions for downstream tasks.

We have opted for the [CLS] token representation when assessing downstream tasks. Additionally, the aforementioned three types of representations are available on https://github.com/a96123155/UTR-LM.

### B.3. Splitting strategies of training and test data for MRL tasks

Sample et al.^7^ employed a read count-based sorting strategy, which we have termed Rank Split, to select 5’ UTRs for training and testing. For the eGFP and mCherry libraries, the top 280,000 and 200,000 sequences were selected, respectively. From each library, the 20,000 UTRs with the highest read counts were allocated for the test set, with the rest reserved for training.

We introduced a method called Random Split to better capture data variability and reflect real-world situations. This method involves randomly selecting 20,000 sequences from the top 280,000 and 200,000 sequences in the eGFP and mCherry libraries, respectively, to form the test set, while the remaining sequences are used for training.

### B.4. Experimental settings of independent test

In the Optimus method^7^, a test set is formed from 100 5’ UTRs of each length that have the deepest read coverage, ensuring approximately 10% of the library. This results in two test sets, each containing 7,600 5’ UTRs—one composed of random sequences and the other of human sequences. The remaining 76,319 random 5’ UTRs are used for model training. Both FramePool and RNA-FM use the same training set as Optimus, but their testing approaches differ. FramePool did not create a validation set and carries out model selection and evaluation using the 7,600 random 5’ UTRs. On the other hand, RNA-FM employs the 7,600 random 5’ UTRs as a validation set to determine optimal parameters and uses the 7,600 real human 5’ UTRs as the test set. For fair comparison, we split the original training set (n = 83,919) into a training set and a validation set at a ratio of 9:1. These sets are used for model training and selection, respectively, with performance then evaluated on the two test sets, each containing 7,600 5’ UTRs.

In the aforementioned experiments, 5’ UTRs with the highest total read counts were used for the test set. Those 5’ UTRs with more reads demonstrated higher resolution than those with fewer reads, and thus more accurately reflected their MRL (see Supplementary Fig. 1^7^). However, prediction methods should also perform well on test sets that are randomly split, as it can be challenging to guarantee high-quality samples in real-world situations. Thus, we conducted 10-fold cross-validation experiments for 83,919 random 5’UTR libraries and 15,555 human 5’ UTR libraries, respectively. The results are displayed in Supplementary Fig. 2.

**Supplementary Fig. 1.**
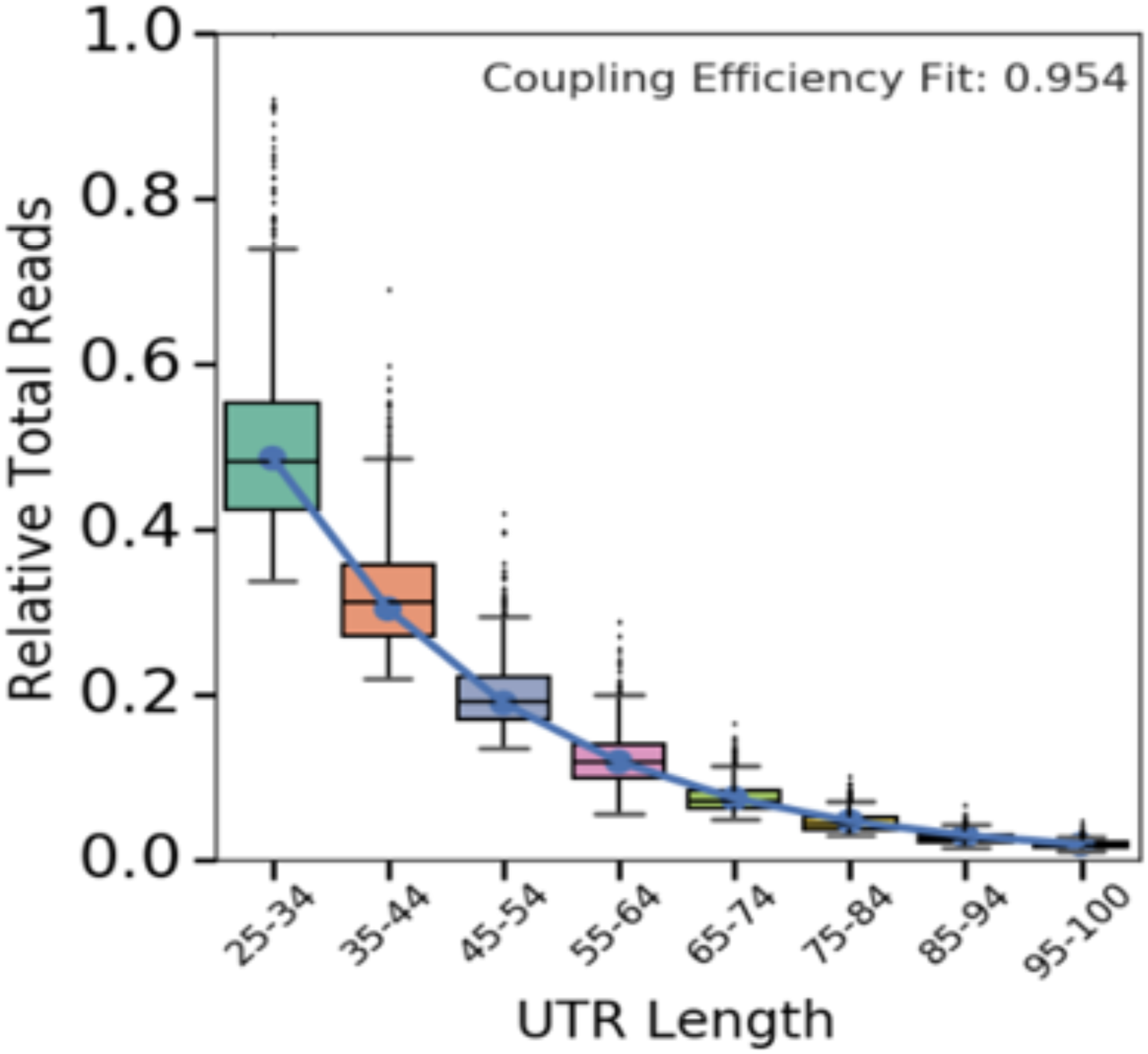
Fit the coupling efficiency on total reads in terms of 5’ UTR length. This figure is proposed by Sample et al.^7^

**Supplementary Fig. 2.**
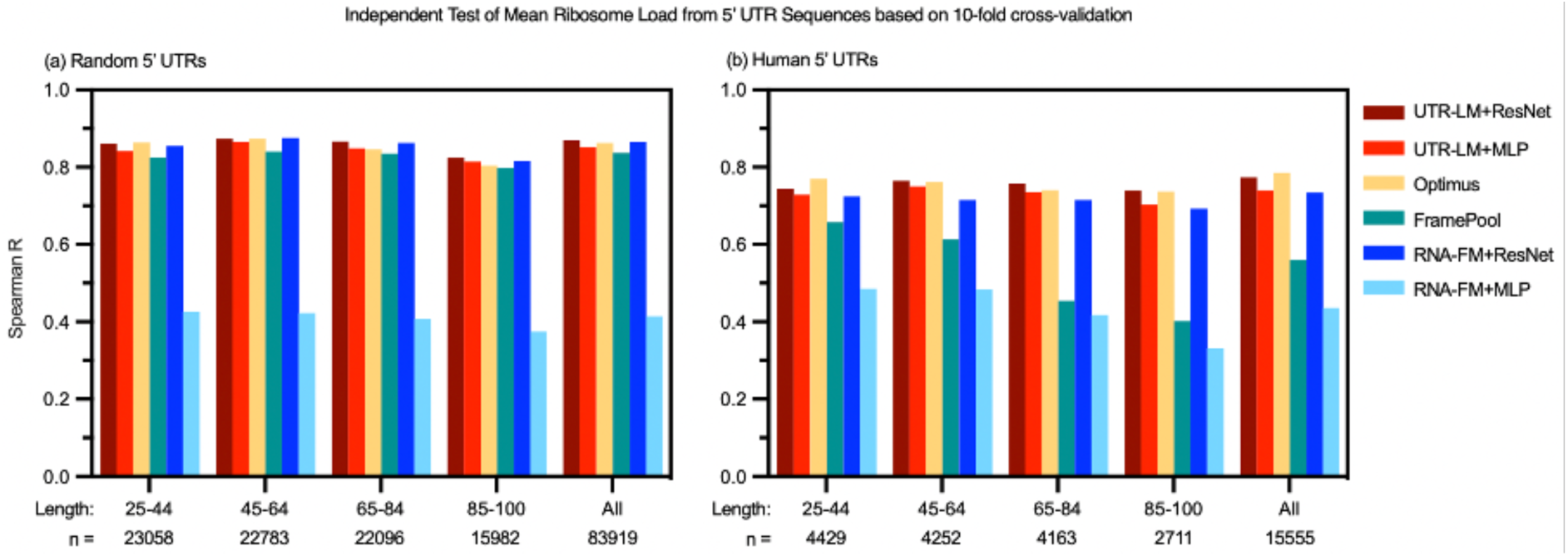
Performance for independent test sets across methods under 10-fold cross-valiation for (a) 83,919 random 5’UTR libraries and (b) 15,555 human 5’ UTR libraries.

## C. Model Interpretation

### C.1. Details of differentiating the five species data

We validated the superiority of the pre-trained embedding by testing its differentiation capacity on data from five species. For the baseline, we used the *k*-mer representation, which is frequently employed as a baseline method in RNA-seq analysis^35^ due to its ability to capture critical features such as secondary structure, splice sites, and binding sites of RNA-binding proteins. Among the various *k*-mer lengths, we selected *k*=4, as it strikes a balance between computational complexity and the amount of information captured. This choice offers advantages over other commonly used methods. For instance, the simpler one-hot encoding may not capture all relevant information in RNA-seq, while the more complex position weight matrices necessitate a larger volume of data to be effective.

## D. Related Work

Recent years have witnessed increasing interest in using language models to decode biological sequences^36^. While there are plenty of protein language models such as ESM-1b^37^, TAPE^38^, ESMFold^39^, and ProGen^40^, there are fewer language models for learning semantic representations of DNA and RNA sequences. Benegas et al.^41^ introduced the Genome pre-trained Network, an unsupervised model learning variant effects, gene structures, and DNA motifs in non-coding DNA. Dalla-Torre et al.^42^ proposed the Nucleotide Transformer, a DNA sequence pre-training method that outperforms specialized methods in multiple prediction tasks using diverse species genomes. Ji et al.^43^ developed DNABERT, pre-trained on the human genome, which effectively predicts promoter regions, splice sites, and transcription factor binding sites after fine-tuning. Yamada et al.^36^ further demonstrated DNABERT’s potential in RNA-related tasks, predicting RNA-binding protein interactions. Akiyama et al.^10^ proposed RNABERT, which effectively utilizes the pre-trained BERT algorithm for non-coding RNA tasks and does not give results for any 5’ UTR application. Chen et al.^9^ introduced RNA-FM, a model pre-trained on 27 million unannotated RNA sequences through self-supervision. Both RNABERT and RNA-FM have demonstrated utilities in structure prediction and function-related tasks. Models like SpliceBERT^44^ and miProBERT^45^ are pre-trained on specific RNA types, such as pre-mRNA and microRNA promoters. To the authors’ best knowledge, no one has attempted to develop a language model specifically for modeling UTR sequences and their functions.

Several studies leverage massively parallel translation assays and specialized machine learning models, like ribosome profiling, to study 5’ UTR variant effects on mRNA translation. Cao et al.^8^ developed random forests for predicting mRNA expression level (EL) and translation efficiency (TE) based on handcrafted features like *k*-mer frequencies, RNA folding energy, 5’ UTR length, and the number of open reading frame. Sample et al.^7^ proposed Optimus, convolutional neural networks (CNN) for predicting the mean ribosome loading (MRL) of 5’ UTRs, extendable to longer human 5’ UTRs. Karollus et al.^6^ modified Optimus using global pooling, known as the FramePool method, to remove length restrictions. Chen et al.^9^ introduced the RNA-FM embedding, which was used in conjunction with a 32-layer residual network (ResNet) to predict MRL. Zheng et al.^5^ developed a multi-task CNN model for TE prediction using multiple data sources.

